# Covalent linkage of the DNA repair template to the CRISPR/Cas9 complex enhances homology-directed repair

**DOI:** 10.1101/218149

**Authors:** Natasa Savic, Femke Ringnalda, Katja Bargsten, Yizhou Li, Christian Berk, Jonathan Hall, Dario Neri, Martin Jinek, Gerald Schwank

## Abstract

The CRISPR/Cas9 targeted nuclease technology allows the insertion of genetic modifications with single base-pair precision. The preference of mammalian cells to repair Cas9-induced DNA double-strand breaks via non-homologous end joining (NHEJ) rather than via homology-directed repair (HDR) however leads to relatively low rates of correctly edited loci. Here we demonstrate that covalently linking the DNA repair template to Cas9 increases the ratio of HDR over NHEJ up to 23-fold, and therefore provides advantages for clinical applications where high-fidelity repair is needed.

## Main text

The CRISPR/Cas9 system is a versatile genome-editing tool that enables the introduction of site-specific genetic modifications^1^. In its most widespread variant a programmable chimeric short guide RNA (sgRNA) directs the Cas9 nuclease to the genomic region of interest, where it generates a site-specific DNA double-strand break (DSB)^2^. In mammalian cells, DSBs are either repaired by non-homologous end joining (NHEJ), or by homology-directed repair (HDR) pathways^3^. While NHEJ is an error-prone process that produces random insertions or deletions (indels)^4^, HDR repairs DSBs accurately from template DNA, and enables the introduction of modifications with single base precision^5^.

Therapeutic applications of CRISPR/Cas9 generally require the precise correction of pathogenic mutations. In mammalian cells, however, DSBs are predominantly repaired by NHEJ. As the thereby induced indels inhibit the CRISPR/Cas9 complex from retargeting the locus, NHEJ directly competes with HDR and reduces precise correction rates. In addition, if the targeted allele is a hypomorph with residual gene function, indels generated by NHEJ could further worsen the clinical phenotype of the disease. In recent years, several attempts have been made to enhance DSB repair by the HDR pathway. These include: i) synchronizing cells in the M phase of the cell cycle prior to CRISPR/Cas9 delivery^6^, ii) limiting Cas9 expression to the S/G2 phase of the cell cycle^7^, iii) chemically modulating the NHEJ and HDR pathways ^8–11^, and iv) rationally designing DNA repair templates with optimal homology arm lengths ^12^. In addition, it has been proposed that the availability of the DNA repair template might present a rate-limiting factor for HDR, and that bringing it in close spatial proximity to the DSBs could therefore enhance HDR editing rates ^13,14^. Based on this hypothesis, we here generated and tested novel CRISPR/Cas variants, in which the DNA repair template is covalently conjugated to Cas9 via ‘click chemistry’ (Fig 1a).

**Figure 1:**
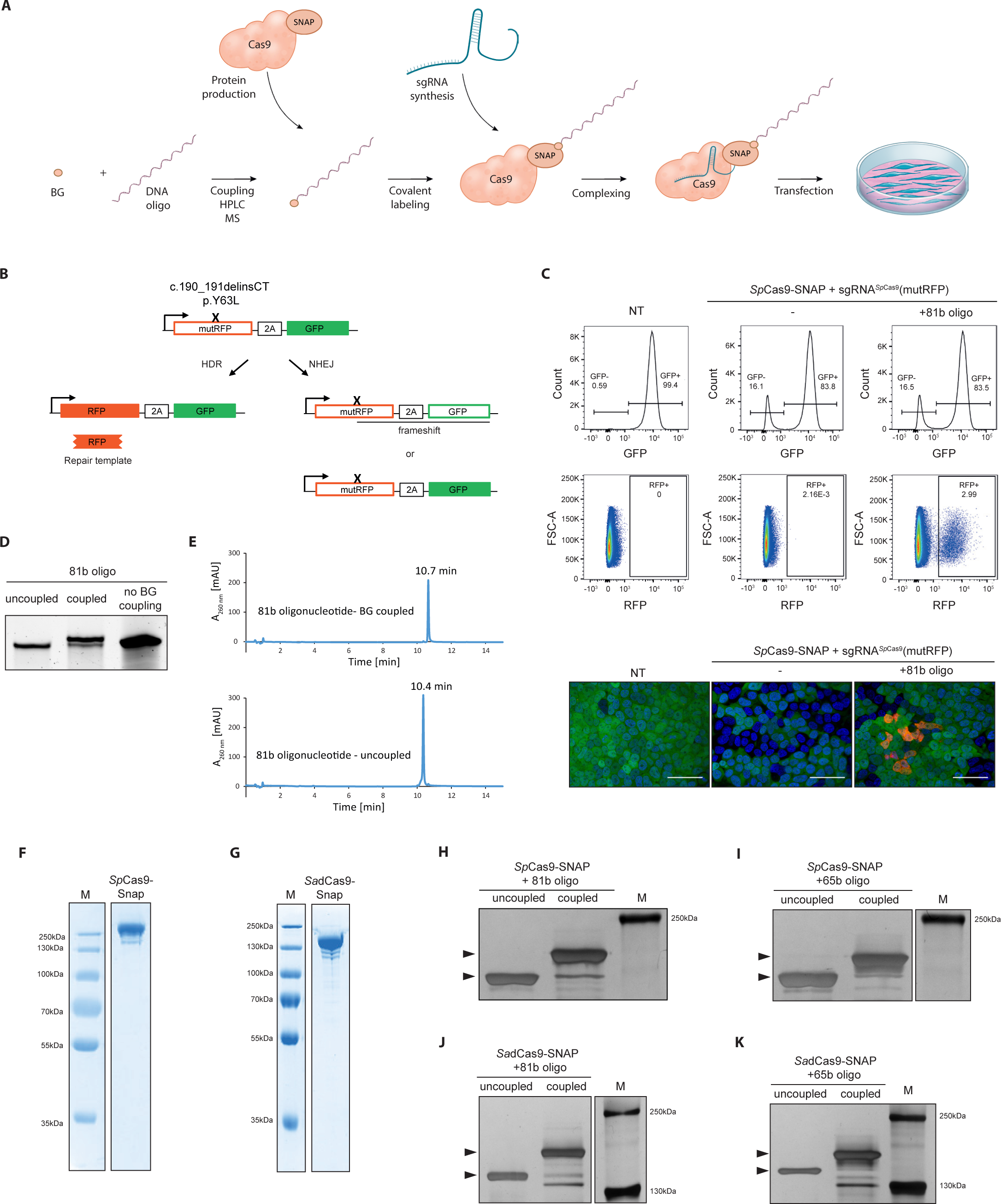
A high-level schematic of `bossDB` platform.

To be able to easily test HDR based editing efficiencies of novel CRISPR/Cas9 variants in a high throughput manner, we first generated a fluorescent reporter system that allowed us to quantify HDR and NHEJ editing frequencies in mammalian cells by FACS (Fig. 1b). In brief, the reporter expresses a green fluorescent protein (GFP), which is preceded by an inactive version of a mutant red fluorescent protein (mutRFP). While precise correction of the mutation via HDR leads to re-activation of RFP activity, the generation of frame shifts via NHEJ leads to loss of GFP activity (Fig. 1b). To test the functionality of the reporter and to determine the optimal length of DNA repair templates, we transfected mammalian cells that stably express a single copy of the reporter with Cas9-sgRNA ribonucleoprotein (RNP) complexes and repair templates of different lengths (Fig. Sup. 1a). In line with previous studies^15^, we found that maximal HDR efficiencies of DSBs are reached with DNA oligonucleotides (oligos) of approximately 80 bases. Nevertheless, as we reasoned that if repair templates are brought in close proximity to DSBs also shorter homology arms could be sufficient, we continued our study with 65-nucleotide (65-mers) and 81-nucleotide (81-mers) DNA repair templates.

In order to link repair oligos to Cas9, we used the SNAP-tag technology, which allows covalent binding of *O*^6^-benzylguanine (BG)-labeled molecules to SNAP-tag fusion proteins ^16^. To generate *O*^6^-benzylguanine (BG)-linked DNA repair templates, we first synthesized amine-modified oligos, and coupled them to commercially available amine-reactive BG building blocks (Fig. 1d, Suppl. Fig. 1b). The BG-linked oligos were further separated from unlinked oligos by HPLC (Fig. 1e), and analyzed by mass spectrometry to confirm purity (Suppl. Fig. 1c). Next, we produced recombinant Cas9 proteins with SNAP-tag fused to the C-terminus (Fig. 1f,g). The fusion proteins were then complexed with the BG-coupled oligonucleotides, and covalent binding was confirmed by SDS-PAGE (Fig. 1 h-k). The protein-oligo conjugate was finally mixed with *in vitro* transcribed sgRNAs targeting the mutRFP locus (Suppl. Fig. 1f,g), generating the Cas9 ribonucleoprotein-DNA (RNPD) complex.

To test if linking the repair template to Cas9 changes the ratio between NHEJ and HDR, we used our reporter system to compare *S. pyogenes* (*Sp*)Cas9 complexes with coupled repair oligos to *Sp*Cas9 complexes with uncoupled repair oligos (Fig. 2a). Notably, the correction efficiency (percentage of HDR in edited cells) with coupled complexes was significantly enhanced, reaching 22% with the 65-mers and 26% with the 81-mers (Fig. 2b, Suppl. Fig. 2a,b). In comparison to uncoupled complexes this represented a 12-and 4-fold increase, respectively (Fig. 2c).

Since it is conceivable that that the RNPD complex could dissociate from the target locus before repair of the DSB is initiated, we designed a two-component system in which the DSB is induced by the *Sp*Cas9 RNP complex, and the repair template is linked to a catalytically inactive *Staphylococcus aureus* (*Sa*)dCas9 that binds in close proximity to the targeted locus. As the inactive *Sa*dCas9 does not induce DSBs, its target sequence is not destroyed, thus avoiding dissociation of the repair template from the locus (Fig. 2d). To test whether this two-component system further enhances gene-repair efficiency, we co-transfected both complexes into the reporter cell line, and quantified HDR and NHEJ rates. Notably, the correction efficiency increased to 30% with 65-mers, and to 33% with 81-mers (Fig. 2e, Suppl. Fig. 2c,d), confirming our hypothesis that longer retention of the repair template at the DSB further enhances HDR rates.

**Figure 2.**
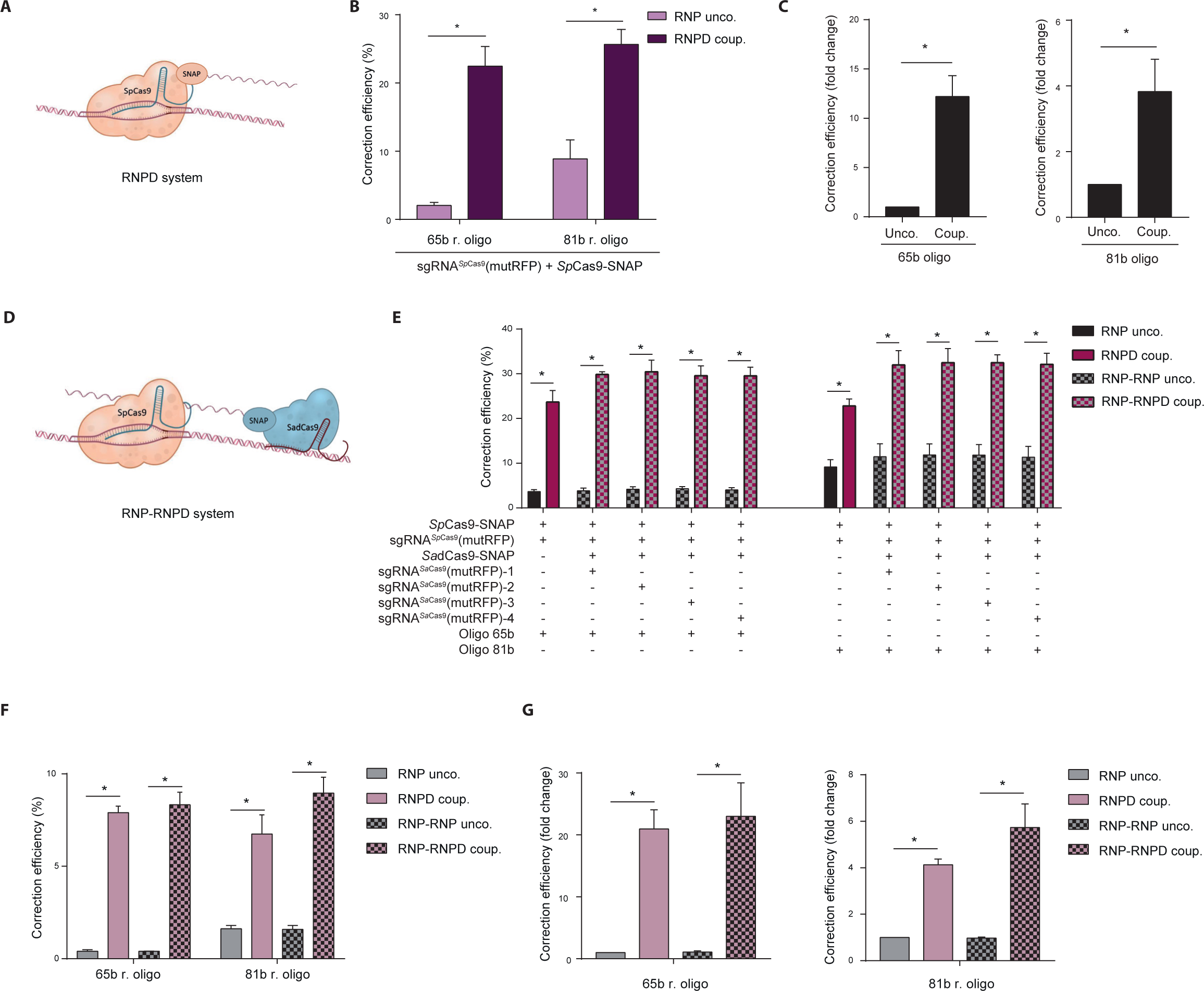
Linking the repair template to the Cas9 RNP complex enhances HDR at the expense of NHEJ. (**a**) Illustration of our novel Cas9 system, in which the repair template is coupled to *Sp*Cas9 (RNPD coup.). (**b,c**) Comparison of the classical Cas9 system (RNP unco.) to our novel system (RNPD coup.). NHEJ and HDR rates were quantified 5 days after transfection by FACS. Results presented as correction efficiency (percentage of HDR in edited cells) (**b**), and as fold change in comparison to the classical Cas9 system (**c**). (**d**) Illustration of the two component system, where the repair template is linked to the catalytically inactive *Sa*dCas9 (RNP-RNPD coup.). (**e**) Comparison of the correction efficiency between the one-component system (a) and the two component system (d). Black and red panels: One component system. Black/grey and red/grey panels: Two component system. RNP-RNP uncoup. (*Sp*Cas9 + *Sa*dCas9 + unlinked repair oligo). RNP-RNPD coup. (*Sp*Cas9 + *Sa*dCas9 with coupled repair oligo). sgRNAs^*Sa*dCas9^(mutRFP)-1-4 bind to different sites in close proximity to the *Sp*Cas9-induced DSB. (**f,g**) Transfection of the one-and two component system at a 5-fold lower concentration. Results presented as correction efficiency (percentage of HDR in edited cells) (**f**), and as fold change in comparison to the classical Cas9 system (**g**). In the two component system sgRNA^*Sa*dCas9^(mutRFP)-3 was used. RNP unco. = (*Sp*Cas9-SNAP-tag + sgRNA^*Sp*Cas9^(mutRFP) + uncoupled oligo.); RNPD coup. = (*Sp*Cas9-SNAP-tag + sgRNA^*Sp*Cas9^(mutRFP) + *Sp*Cas9-SNAPtag + coupled oligo); RNP-RNP unco. = (*Sp*Cas9-SNAPtag + sgRNA^*Sp*Cas9^(mutRFP) + *Sa*dCas9-SNAP-tag, sgRNA^*Sa*dCas9^(mutRFP) + uncoupled oligo); RNP-RNPD coup. = (*Sp*Cas9-SNAP-tag + sgRNA^*Sp*Cas9^(mutRFP) + *Sa*dCas9-SNAP-tag, sgRNA^*Sa*dCas9^(mutRFP) + *Sa*dCas9-SNAP-coupled oligo). All values are shown as mean±s.e.m of biological replicates; *P<0.04 with n=3 (**f,g**) and n=4 (**b,c,e**) (n represents the number of biological replicates). A one-tailed Mann-Whitney test was used for comparisons.

*In vivo,* the delivery efficiency of RNPs and oligos is generally lower than *in vitro*. Thus, if the repair template is not bound to Cas9, there is substantial probability that only one of the two components is delivered into the cell. In addition, at lower transfection efficiencies fewer repair templates are present in the nucleus and in close proximity to the targeted locus, potentially decreasing HDR rates. As we presumed that linking the repair oligo to Cas9 should largely overcome these issues, we investigated whether the repair efficiency with RNPD complexes is affected when transfected at 5-fold lower concentrations. Importantly, although under these conditions the correction efficiencies were generally lower with both bound- and unbound Cas9 complexes, the difference between the two systems was even more pronounced. Compared to the uncoupled RNP complex, the RNPD system yielded a 21-fold and a 4-fold increase in repair efficiency with 65-mer and 81-mer repair template oligos, respectively. Similarly, the two-component RNP-RNPD system led to a 23-fold increase with 65-mers and a 6-fold increase with 81-mers (Fig. 2f,g, Suppl. Fig. 2e,f). Our results suggest that linking the repair template to the Cas9 complex leads to improved correction efficiency compared to the classical CRISPR/Cas system, and that this effect is even more pronounced when CRISPR/Cas components are delivered at lower concentrations.

Direct delivery of Cas9 RNP complexes into tissues promises great potential for therapeutic applications. Compared to genetically encoded systems, RNPs avoid the danger of genomic integration, and have fewer off-target effects due to their limited lifetime. The method presented here enhances HDR-based gene editing without altering endogenous cellular processes, and is thus poised to further drive the CRISPR/Cas technology towards clinical translation.

## Methods

Please see Supplementary Tables 1–4 for a list of the DNA sequences used in this manuscript.

### Plasmids

All plasmids used in this study are listed in Supplementary Table 4.

Cloning of pNS19-LV-mutRFP-2A-GFP: pEGIP (addgene plasmid #26777) was mutagenized using QuikChange Lightning Multi Site-Directed Mutagenesis Kit (Agilent Technologies) to destroy the start codon of eGFP. Next the vector was linearized with BamHI and In-Fusion HD Cloning Plus CE (Takara) was used to insert the mutRFP-2A gBlocks Gene Fragment (Integrated DNA Technologies).

Cloning of pNS20-SpCas9-SNAP: pMJ922-SpyCas9-GFP bacterial expression vector was a kind gift from Prof. Martin Jinek. GFP was digested using BamHI and KpnI, and SNAPtag-NLS gBlocks (Integrated DNA Technologies) were integrated using In-Fusion HD Cloning Plus CE (Takara).

Cloning of pNS38-SadCas9-SNAP: pAD-SaCas9-GFP was generated by replacing the SpCas9 coding sequence in pMJ922 with SaCas9 sequence using Gibson cloning ^16^. QuikChange Lightning Multi Site-Directed Mutagenesis Kit (Agilent Technologies) was used to remove the stop codon and to introduce the D10A and N580A mutations into the *Sa*Cas9 gene. Subsequently, GFP was cut out using BamHI and KpnI, and replaced by a SNAP-tag-NLS gBlock (Integrated DNA Technologies) using In-Fusion HD Cloning Plus CE (Takara).

### Benzylguanine coupling reaction

Synthetic oligonucleotides with a 5′-Amino Modifier C6 functional group (100µM) (Integrated DNA Technologies) were incubated with benzylguanine-GLA-NHS (1 mM) (NEB) and Hepes pH8.5 (200mM) for 60 minutes at 30 °C. Coupling reactions were performed in following ratios: 30:1, 60:1 and 100:1 BG-GLA-NHS: amino modified oligo. After the coupling reaction all oligos were purified by ethanol precipitation. Repair oligo sequences can be found in Supplementary Table 3.

### Denaturating PAGE

The benzylguanine (BG) coupled reactions were run on 20% polyacrylamide TBE gel containing 8M urea at 200V for 60 minutes. The gel was stained for 30 minutes in 1x TBE containing Sybr^®^Gold (Invitrogen), and imaged with a UV transilluminator (Biorad).

### HPLC purification and MS analysis of the repair oligos

Benzylguanine coupled oligos were purified on an Agilent 1200 series preparative HPLC fitted with a Waters XBridge Oligonucleotide BEH C18 column, 10 × 50mm, 2.5µm at 65°C using a gradient of 5-25% buffer B over 8 min, flow rate= 5 ml min-1. Buffer A was 0.1M triethylammonium acetate, pH 8.0. Buffer B was methanol. Fractions were pooled, dried in a speedvac and dissolved in H2O. Analysis of the purified BG-oligonucleotide was conducted on an Agilent 1200/6130 LC-MS system fitted with a Waters Acquity UPLC OST C18 column (2.1×50 mm, 1.7 µm) at 65°C, with a gradient of 5-35% buffer B in 14 min with a flowrate of 0.3 mLmin−1. Buffer A was aqueous hexafluoroisopropanol (0.4M) containing triethylamine (15 mM). Buffer B was methanol.

### Expression and purification of Cas9-SNAP

Snap-tagged *Streptococcus pyogenes* Cas9(SpCas9-SNAP) and *Staphylococcus aureus* dCas9 (SadCas9-SNAP) proteins were expressed in *Escherichia coli* BL21 (DE3) Rosetta 2 (Novagen) fused to an N-terminal fusion protein containing a hexahistidine affinity tag, the maltose binding protein (MBP) polypeptide sequence, and the tobacco etch virus (TEV) protease cleavage site. The cells were lysed in 20 mM Tris pH 8.0, 500 mM NaCl, 5 mM Imidazole pH 8.0. Clarified lysate was applied to a 10 ml Ni-NTA (Qiagen) affinity chromatography column. The column was washed by increasing the imidazole concentration to 10 mM and bound protein was eluted in 20 mM Tris pH 8.0, 250 mM NaCl, 100 mM Imidazole pH 8.0. To remove the His_6_-MBP affinity tag, the eluted protein was incubated overnight in the presence of TEV protease. The cleaved protein was further applied to a heparin column (HiTrap Heparin HP, GE Healthcare) and eluted with a linear gradient of 0.1-1.0 KCl. The was further purified by size exclusion chromatography using a Superdex 200 16/600 (GE Healthcare) equilibrated in 20 mM HEPES pH 7.5, 500 mM KCl.

### Covalent binding of Cas9-SNAP protein and BG-coupled oligonucleotide

Repair oligo templates coupled to BG were incubated with Cas9-SNAP proteins on the same day when the transfection is performed. BG-coupled oligos (2.2 pmols) were mixed with either SpCas9-SNAP or SadCas9-SNAP (2.2 pmols) and incubated for 60 minutes at 30°C.

### SDS-PAGE gels

For confirming successful labeling of the Cas9-SNAP proteins with the BG-coupled oligonucleotides, BG-coupled and uncoupled oligonucleotides were mixed with either SpCas9-SNAP, SadCas9-SNAP or only the Cas9-SNAP proteins alone, reactions were incubated for one hour at 30°C. For the SNAP-Vista^®^Green (NEB) substrate, the protein was incubated for 30 minutes on 30°C in the dark. After incubation, reactions (300ng) were loaded on 6% SDS-PAGE gel and run at 80V for 160 minutes. Gels that were containing BG-Vista Green (NEB, SNAP-Vista^®^Green), were imaged prior to silver staining. The green fluorescence signal of the SNAP-tag was detected with a UV transilluminator (Biorad). Subsequently, silver staining was completed using the Pierce™ Silver Stain Kit (Thermo Scientific) according to manufacturer instructions, and imaged with a UV transilluminator (Biorad).

### Production of sgRNAs

sgRNAs were generated from DNA templates using the T7 RNA Polymerase (Roche) *in vitro* transcription (IVT) kit. In short, sgRNA specific primers that also contain the T7 sequence were annealed with a common reverse primer that contains the sequence of the sgRNA scaffold (final concentrations 10µM). DNA was purified with the QIAquick purification (Qiagen) kit and eluted in DEPC-treated water. PCR products were run on agarose to estimate concentration and to confirm amplicon size. *In vitro* transcription was performed at 37°C overnight. For purification, DNase I was added to the sgRNAs and incubated for 15 minutes at 37°C, and subsequently ethanol precipitation was performed overnight at −20°C. The sgRNAs were then further purified using RNA Clean & Concentrators (Zymo Research). Before use, all sgRNAs were checked on denaturing 2% MOPS gels. Complete sequences for all sgRNA protospacers and IVT primers can be found in Supplementary Table 1 and 2, respectively.

### Lentivirus production

HEK293T were PEI transfected with following plasmids: pNS19-LV-mutRFP-2A-eGFP, Pax2 and VSV-G. After 12 hours, the supernatant was discarded and changed to DMEM plus 10% FBS. 24 and 72 hours post-transfection, the media was collected and filtered through 0.45 µm filter and centrifuged at 20 000 G for 2:00 hours at 4°C. The pellet was then resuspended in 1 ml of DMEM and stored at −80°C.

### Fluorescent reporter generation

HEK293T cells were transduced with a lentiviral vector carrying the fluorescent reporter construct. Serial virus dilutions were used to isolate clonal populations using Puromycine selection (2µg/ml) for 2 weeks.

### Cell culture and reagents

HEK293T cells were obtained from ATCC and verified mycoplasma free (GATC Biotech). The HEK293T reporter line was maintained in DMEM with GlutaMax (Gibco). Media was supplemented with 10% FBS (Sigma), and 100 µg/mL Penicillin-Streptomycin (Gibco). Cells were passaged three times per week. Cells were grown at 37°C in a humidified 5% CO^2^ environment.

### Transfection reactions

HEK293T reporter cells were seeded in 24-well plates at 120.000 – 140.000 cells per well, 1 day prior to transfection. On the day of transfection, RNP and RNPD complexes (2.2 pmols) were complexed with sgRNA (3.88 pmols) in Opti-MEM (Invitrogen) and briefly vortexed, followed by adding 3 µl the Lipofectamine^®^2000 reagent (Invitrogen) with Opti-MEM. The resulting mixture was incubated for 15 min at room temperature to allow lipid particle formation. After 15 minutes of incubation at room temperature, the mixture was dropped slowly into the well. One day post-transfection, cells were transferred to an 10cm dish.

### Flow cytometry analysis

For flow cytometry analysis, HEK293T reporter cells were analysed 5 days after transfection. Cells were trypsinized with TrypLE™Express Enzym (Gibco), and resuspended in FACS buffer containing PBS/1% FBS/1% EDTA. Sytox ™Red was added for the exclusion of dead cells. Data were acquired on a BD LSR Fortessa cell analyser (Becton-Dickinson) and were further analysed using FlowJo software (FlowJo 10.2). In all experiments, a minimum of 200.000 cells were analysed. Gating strategy: Forward versus side scatter (FSC-A vs SSC-A) gating was used to identify cells of interest. Doublets were excluded using the forward scatter height versus forward scatter area density plot (FSC-H vs. FSC-A). Live cells were gated based on Sytox-Red-negative staining. Live-gated cells were further used to quantify the percentage of eGFP negative and turboRFP positive populations.

### Fluorescence microscopy

HEK293T reporter cells were imaged 7 days after transfection. Transfected cells were grown on Poly-L-lysine coated 8-well glass chamber slides (Vitaris) to 80-90% confluence. Hoechst 33342 (Thermo Scientific, Pierce™) was added in the cell culture media to a final concentration of 0.1 µg/ml, and cells were incubated for 10 min at 37 °C, 5% CO_2_, prior to the image session. Confocal imaging was performed using a Leica DMI8-CS (ScopeM) with a sCMOS camera (Hamamatsu Orca Flash 4.0). The laser unit for confocal acquisition (AOBS system) contains 458, 477, 488, 496, 514nm (Argon laser), 405nm, 561nm, 633nm. Images were acquired using Leica LAS X SP8 Version 1.0 software, through using a 20x 0.75NA HC PLAN APO CS2 objective. Imaging conditions and intensity scales were matched for images presented together. Images were analysed using the Leica LAS AF (Lite) software version 3.3. Confocal images were processed using ImageJ software (Version 1.51n).

### Statistical analyses

Statistical analyses were conducted using Graphpad’s Prism7 software. A Mann-Whitney test was conducted for two-sample analyses (*P < 0.04). All values are shown as mean ± s.e.m of biological replicates.

### Data availability

The data that support the findings of this study are available within the paper and its Supplementary Information.

